# Validity of Optical Heart Rate Measurement in Commercially Available Wearable Fitness Tracking Devices

**DOI:** 10.1101/2022.09.29.510075

**Authors:** Jason Thomas, Patrick Doyle, J. Andrew Doyle

**Affiliations:** Department of Kinesiology and Health, College of Education and Human Development, Georgia State University, Atlanta, Georgia, United States of America

**Keywords:** photoplethysmography, heart rate monitor, smartwatch, fitness tracking device

## Abstract

**Background:** Wearable fitness tracking devices have risen in popularity for athletes and the general population and are increasingly integrated into smartwatch technology. Many devices incorporate optical heart rate (HR) measurement by photoplethysmography which provides data used to monitor and track exercise training intensities, progress, and other health and fitness related parameters.

**Objective:** To determine the validity of optical heart rate measurement in three fitness tracking devices while resting, walking, and running.

**Methods:** **T**wenty subjects (10 male, 10 female) completed the research study based on the ANSI/CTI standards for physical activity monitoring of heart rate under 4 different conditions: sedentary (SED), treadmill walking (WLK), running (RUN), and dynamic running/walking (DYN). Subjects wore 3 optical heart rate devices: Polar OH1 (OH1) on the right forearm, Apple Watch 4 (AW4) on the right wrist and Garmin Forerunner 945 (FR945) on the left wrist. A Polar H10 (H10), a chest strap device, was the criterion HR measurement device. SED, WLK, and RUN were all 7-minute protocols with 1 minute of standing, 5 minutes of prescribed activity, and 1 final minute of standing. The DYN protocol was a 12-minute protocol with 1 minute of standing, 10 minutes of variable intensity walking and running, and 1 minute of standing. Raw HR data was extracted from each device and temporally aligned with the criterion H10 HR data for analysis.

**Results:** The mean absolute deviation (MAD, measured in beats per minute) for the three experimental devices (OH1, AW4, FR945, respectively) for SED was 1.31, 1.33, and 2.03; for WLK was 2.79, 2.58, and 5.19; for RUN were 4.00, 4.29, and 6.51; and for DYN was 2.60, 2.44, and 2.44. The mean absolute percent error (MAPE) for the three experimental devices (OH1, AW4, FR945, respectively) for SED was 1.78%, 1.89%, and 2.81%; for WLK was 3.15%, 3.18%, and 5.93%; for RUN was 3.43%, 3.51%, and 5.25%; and for DYN was 2.05%, 1.95%, and 5.47%. The intraclass correlation for each device across all conditions was .991 (OH1), .984 (AW4), and .697 (FR945).

**Conclusions:** At rest, and during both steady-state and variable-speed treadmill walking and running, the Polar OH1, Garmin Forerunner 945, and Apple Watch 4 optical HR monitors demonstrated a level of accuracy well within that required by the ANSI/CTA Standard (2018) for physical activity monitoring devices for heart rate measurement (i.e., <10% Mean Absolute Percent Error). Therefore, consumers can have confidence that these devices provide HR data with accuracy that conforms to the performance criteria recommended for consumer electronics.

## Introduction

### Background

Wearable fitness tracking devices have risen in popularity over the past decade and have been the top fitness trend numerous years while approaching nearly a $100 billion industry (1). These devices were initially developed as either rudimentary mechanical pedometers attached to a shoe or waist band, or electrode chest strap heart rate monitors that are often deemed uncomfortable and cumbersome. As technology has advanced, wearable fitness devices have integrated improved technologies including GPS, accelerometers, altimeters, and photosensors. Further, they are increasingly integrated into more user-friendly and comfortable devices, specifically wrist-worn watches, and arm bands. As heart rate (HR) monitoring is arguably the key component of fitness monitoring, a principle technological advance has been the integration of photoplethysmography (PPG), which uses a light emitting diode and photosensor to measure microvascular blood volume changes which is consequently associated with heart rate (2).

The advancement and integration of PPG technology into wrist-worn devices has granted the end-user with a wealth of information including caloric expenditure, oxygen consumption (VO_2_), heart rate variability, sleep patterns, recovery, and training intensity. All this information provided to users is based on manufacturer-specific algorithms computed from heart rate collected via PPG technology. Therefore, the validity of the heart rate measurement from these PPG devices is of key importance.

Several studies have been completed to assess the validity of a variety of activity tracking devices which use the PPG technology. Although several devices, including the OH1, Apple Watch series, and Garmin Forerunner series, have been deemed valid, the results of the studies must be interpreted narrowly as various methodological differences or concerns exist between studies. As device availability has grown immensely and rapidly, the current body of research lacks results that can more confidently discern the validity of the devices across the general population.

The OH1 has been previously studied and was deemed valid for moderate and high intensity activities (3, 4), appeared more valid compared to a wrist-based device by the same manufacturer(5), and showed decreased validity with arm-based activities (i.e. tennis) (6). A key limitation of these studies is the application of the study results to the wider population as the studies lacked balanced diversity in either BMI, skin tone, or sex. Likewise, studies using the Apple Watch series have suggested device validity, but different methodological issues exist. The methodological concerns were comprised of various issues such as recording heart rate from a single timepoint (7), using a model of tachycardia (8), implementing a single subject design (9), or failing to report key validity metrics such as MAD, MAPE, and ICC (10). Additionally, these studies also lacked the diversity in key subject demographics, similar to the limitations with OH1 research. The Garmin Forerunner series has a very limited amount of information available in the literature. The existing data has either suggested poor validity in prior versions to the FR945 (7) or has suffered from methodological issues related to heart rate recording frequency (11).

Aside from specific device validity, the current body of research for all PPG activity tracking devices suffers from numerous methodological differences that limits our ability to apply the results to the general population. Existing studies generally lack cohesion between different exercise types, intensities, and durations. Some studies have been completed to assess the validity of a single device across multiple exercise modalities (4), while others have researched numerous devices across a variety of intensities and exercise modalities, but with shorter data recording times (12). More recent studies have investigated multiple devices, but intensity was extremely high and duration extremely short (13). Other studies have utilized different modes of exercise but lacked varying levels of intensity within the modes (14, 15). A recent study implemented activity modes and intensity with a better variation, but only incorporated a single demographic (Caucasian) in the subject group (10)

The lack of variation in subject demographics is visible across many studies. Variations in skin tone, BMI, sex, and age have been suggested as potential confounding factors to proper validity testing for PPG technology. Variations in skin tone appears to affect validity as use of a typical green-light LED diode (often integrated into many devices) has resulted in a 1.04 BPM error rate in light-skinned individuals, and as much as a 10.9 BPM error in dark-skinned individuals (16). There is also evidence to suggest that as BMI increases, PPG waveform can change as much as 43% between obese and non-obese individuals (17). Some studies have attempted to address these concerns but have had limitations. For instance, a recent study did investigate skin tone and PPG using an Apple Watch, but the study subjects only represented 3 of the 6 Fitzpatrick skin tone designations (18). Another study had all 6 skin tones represented, but only had 10 subjects total such that certain skin tones were only represented by a single subject (19). Additionally, few studies have specifically recruited subjects to represent a diversity in BMI or gender.

Recently, the American National Standards Institute (ANSI) and Consumer Technology Association (CTA) developed the ANSI/CTA standards for investigating the validity and reliability of consumer electronic fitness devices. These standards provide a consistent, balanced, and equitable basis for subject selection and activity parameters so that consumer devices can be evaluated in a standardized manner. The activity parameters outline optimal intensity levels and duration for different modes of activity. Additionally, subject selection requirements ensure a diverse population relative to age, gender, body mass, and skin tone or complexion.

### Study Objective

PPG technology is being widely implemented to determine HR in an increasing number of devices to appeal to a broader market of consumers globally. As such, it is important to determine if the existing device validity evidenced by previous studies is representative of a diverse population and activities or if the results can only be applied to the limited subject demographics and activities of the respective studies. Therefore, the purpose of this study is to evaluate the heart rate measurement validity of three consumer photoplethysmographic heart rate monitors compared to an accepted criterion device in accordance with current standards of ANSI/CTI.

## Methods

### Participants

Twenty healthy subjects (10 males and 10 females) voluntarily completed the study. Subject characteristics are presented in Table 2. All subjects were educated on the risks of the procedures and gave informed consent prior to the start of the protocols. Subjects were recruited verbally from faculty, staff, and students within the university or by e-mail through a local running club. The study was approved by the Institutional Review Board of Georgia State University.

**Table 1.** Treadmill Intensities

|  | Intensity |  |  |  |
| --- | --- | --- | --- | --- |
| <u>Testing Condition</u> | <u>Moderate</u> | <u>High</u> | <u>Very High</u> | <u>Elite</u> |
| 2 - Steady State Walk | 2.5 MPH | 2.7 MPH | 3.0 MPH | 3.3 MPH |
| 3 - Steady State Run | 5.0 MPH | 6 MPH | 7.0 MPH | 8.0 MPH |
| 4 - Dynamic<br>(Run/Faster/Fastest) | 5/5.5/6.0 MPH | 6/6.7/7.3 MPH | 7/7.7/8.3 MPH | 8/8.7/9.3 MPH |

**Table 2.** Subject characteristics

| <u>Sex</u> | <u>BMI</u><br><u>(kg/m<sup>2</sup>)</u> | <u>Fitzpatrick</u><br><u>Score 1-3</u><br><u>(n)</u> | <u>Fitzpatrick</u><br><u>Score 4-6</u><br><u>(n)</u> | <u>Height</u><br><u>(m)</u> | <u>Weight</u><br><u>(kg)</u> | <u>Body Fat</u><br><u>(%)</u> |
| --- | --- | --- | --- | --- | --- | --- |
| Male (n = 10) | 24.86 | 6 | 4 | 1.73 | 74.54 | 13.23 |
| Female (n = 10) | 23.72 | 8 | 2 | 1.64 | 63.62 | 28.01 |
| All Subjects | 24.3 | 14 | 6 | 1.7 | 69.1 | 20.6 |

### Devices

Four heart rate measurement devices, three experimental and one criterion device, were used for this study. All devices were updated with the most recent software and firmware prior to the start of the study. No further updates were installed on the devices during the data collection period so that firmware and software remained consistent throughout the study. The criterion device was the Polar H10 (H10; Firmware 3.0.50, Polar Electro, Kempele, Finland), an electrode chest-strap heart rate monitor. The Polar H10 uses existing technology from its predecessor Polar H7 which has been validated as above 99% accurate compared to ECG in previous studies (20). The three PPG experimental devices were the Polar OH 1 (OH1; Firmware 2.0.10, Polar Electro, Kempele, Finland), Apple Watch 4 (AW4; Watch OS 5.3.2, Apple, Inc., Cupertino, California) and Garmin Forerunner 945 (FR945; Firmware 2.80, Garmin Ltd., Schaffhausen, Switzerland). The device placement locations were consistent between subjects with OH1 located on the right anterior forearm, AW4 on the right wrist, GF945 on the left wrist. The H10 was fitted on the anterior thorax at the level of the xiphoid process with conduction gel to ensure signal transmission.

### Procedures

Data collection for each subject was completed in a single session and devices were not moved from their specific placement location throughout the entirety of the session. Subjects arrived at the Applied Physiology Laboratory at Georgia State University or the headquarters of a local running club according to their preferred location. After subjects completed informed consent, investigators recorded anthropometric information including subject-described Fitzpatrick score for skin tone, body mass via calibrated digital scale, body fat percentage via 3-site skinfold test, age, and sex. Subjects were then verbally informed of the study protocol, which was a running and walking protocol completed on Woodway treadmills, a Pro XL at the university laboratory and a Desmo S at the local running club (Woodway USA, Inc., Waukesha, WI). Subjects reported a general training intensity level (intensity) described as moderate, high, very high, or elite intensity based upon personal preference and abilities. Walking and running intensities were then assigned by investigators based on this information. Details about the intensity levels are depicted in Table 1.

### Testing Conditions

For each subject, data collection was completed for all 4 testing conditions in a single session. Each testing condition included 1 minute of quiet sitting both prior to and after the treadmill protocol. SED, WLK, and RUN were 7 minutes in length, including the quiet sitting. DYN was 12 minutes in length, including the quiet sitting. For SED, subjects remain seated and motionless for 5 minutes. For the WLK and RUN, subjects completed 5 minutes of activity at the assigned treadmill speed intensity, which investigators set manually for each trial. For DYN, a time-based running and walking protocol, each treadmill was identically pre-programmed with 4 different programs to adjust speed at specific time intervals according to the assigned intensity as seen in Table 1. Walking speed during DYN matched the same intensity speed as WLK condition. Between each testing condition, subjects rested for 5 minutes to allow heart rate to return to normal.

### Data Acquisition

Data from the OH1 and H10 were transmitted from the device via Bluetooth to an iPad Mini running the Performtek app (Valencell, Inc. Raleigh, North Carolina). The Performtek app allows for connection of multiple devices and records device data, including heart rate, for side-by-side comparison. Data from the AW4 was downloaded to RunGap software (CTRL-N ApS, Skødstrup, Denmark) which was then converted to .csv format and imported to Excel. The AW 4 could not be adjusted to record at a specific frequency and required manual data alignment with the same time points of the criterion device for proper analysis. The GF945 data were downloaded as a raw data file (.tcx file) via device sync with Garmin Connect. The H10, OH1, and GF 945 were all programmed to record heart rate at 1 Hz. OH1, H10, and GF945 data were then converted to .csv and imported into a Microsoft Excel (Microsoft Corporation, Redmond Washington) spreadsheet for analysis.

### Statistical Analysis

After being organized in Excel, data were imported into SPSS 27 (SPSS; IBM Corporation, Armonk, NY) for further analysis. Mean Absolute Difference (MAD) and Mean Absolute Percent Error (MAPE) were calculated for each device for each protocol in Excel. T-tests for the difference between experimental device and criterion device for each stage of each protocol were conducted in SPSS to determine mean difference and standard deviation. Pearson’s R correlation and intraclass correlation (ICC) were calculated to determine general correlation between devices and absolute agreement between devices, respectively. Lastly, Bland-Altman plots were created with mean bias and upper and lower limits of agreement. ANSI/CTA standards deem any device with a MAPE ≤ 10% as valid.

## Results

### Subject Characteristics

Basic subject characteristics are presented in Table 2. Recruitment of the subject population was coordinated to adhere to the ANSI/CTA standards for device research such that the minimum percentages of subjects met criteria for Body Mass Index (BMI), Fitzpatrick Score (i.e., skin tone), and sex. The standards as of the ANSI/CTI-2065 were (over the age of 18), sex (no less than 40% male/female), skin tone (minimum 25% from lighter scale and minimum 25% from darker scale), and body mass (minimum 10% above 25 kg/m^2^ and minimum 10% below 20 kg/m^2^). Additionally, a minimum of 20 subjects is recommended.

### General Device Results

Results for all devices can be seen in Tables 3 and 4. More detailed device results based on specific test conditions can be seen in Appendix 1. Both MAD and MAPE are device HR to criterion HR comparisons for all subjects during the entire 7 or 12 minutes of each testing condition. The 7-minute testing conditions had approximately 420 data points (HR measurements) per subject and the 12-minute testing conditions had approximately 720 data points per subject. As the AW4 did not allow for 1Hz HR recording, data points were fewer resulting in approximately 220 data points per subject for the 7-minute protocols and approximately 365 data points per subject for the 12-minute protocol. MAPE must be ≤ 10% to be considered valid according to ANSI/CTA-2065 standards. Using this this threshold, each device was considered valid for each condition tested, although the devices did produce differing results for both MAD and MAPE. Bland-Altman plots for each device’s data aggregated across all conditions can be seen for AW4, FR945, and OH1 in Figures 1, 2, and 3, respectively. Device by test condition Bland-Altman plots can be seen in Appendix 1.

**Fig 1.**
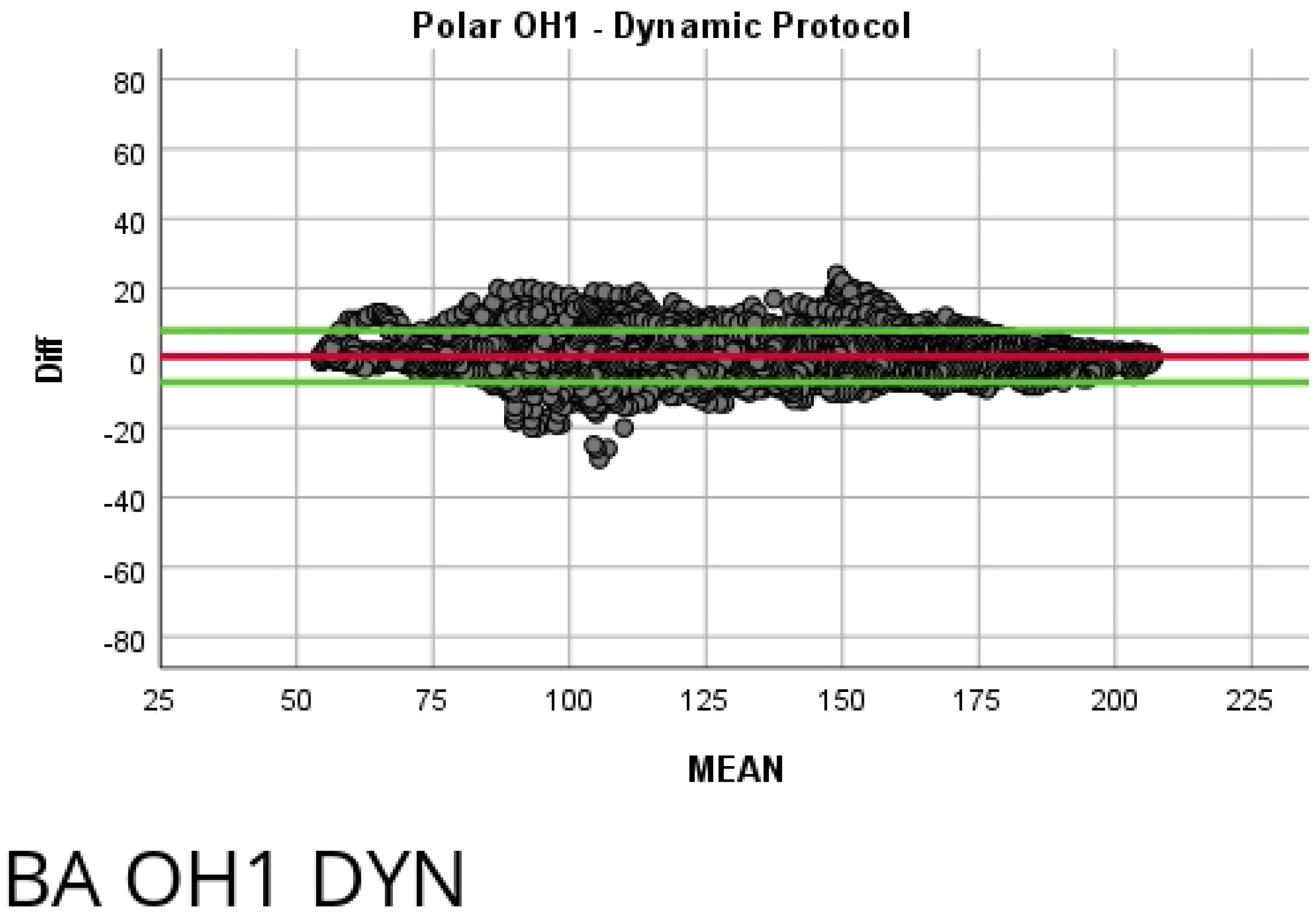
Bland-Altman Plot of all protocols for Polar OH1. Mean bias of 0.59 with upper and lower limits of agreement of 10.586 and −9.406, respectively.

**Fig 2.**
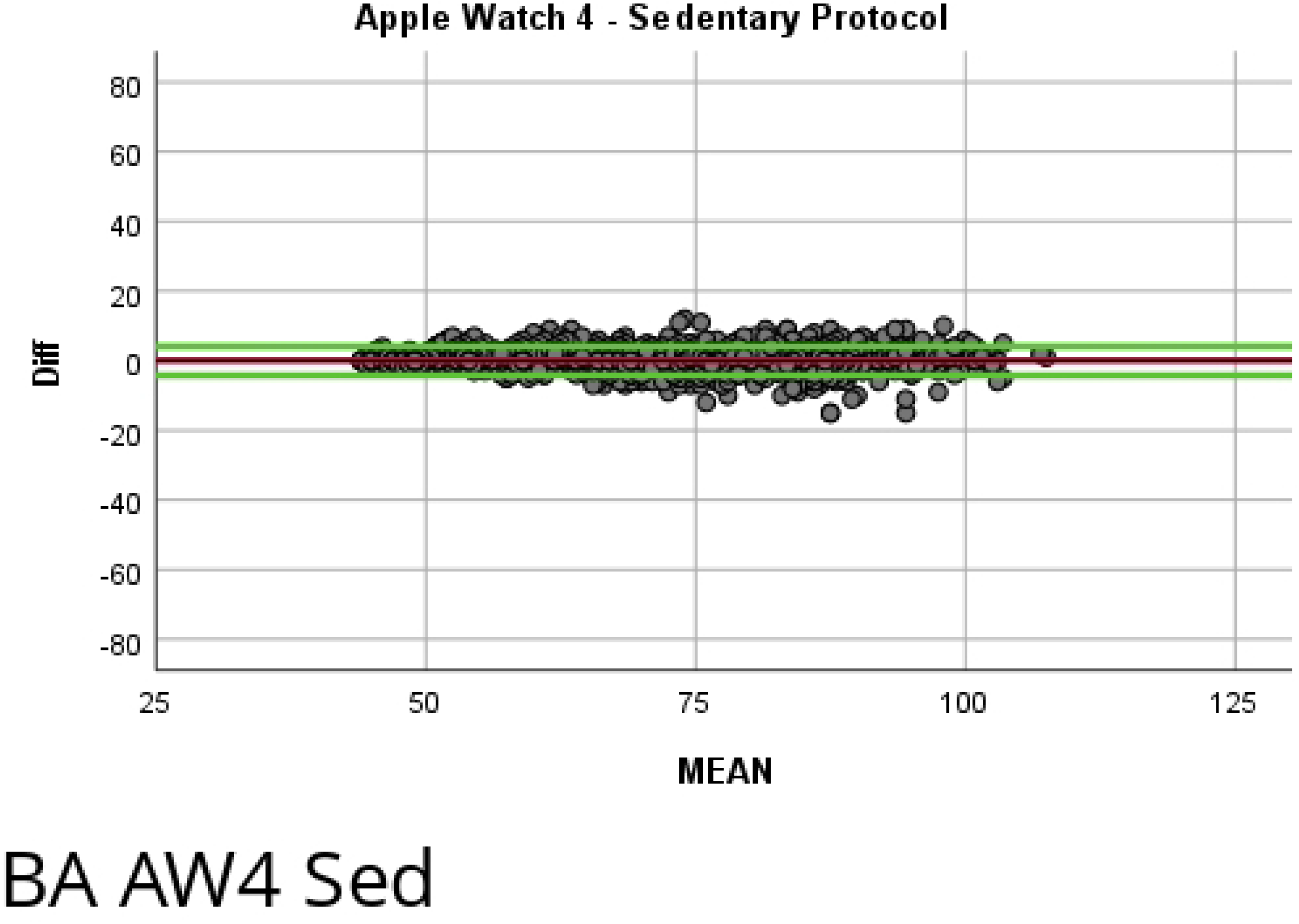
Bland-Altman Plot of all protocols for Apple Watch 4. Mean bias of 0.33 with upper and lower limits of agreement of 13.974 and −13.314, respectively.

**Fig 3.**
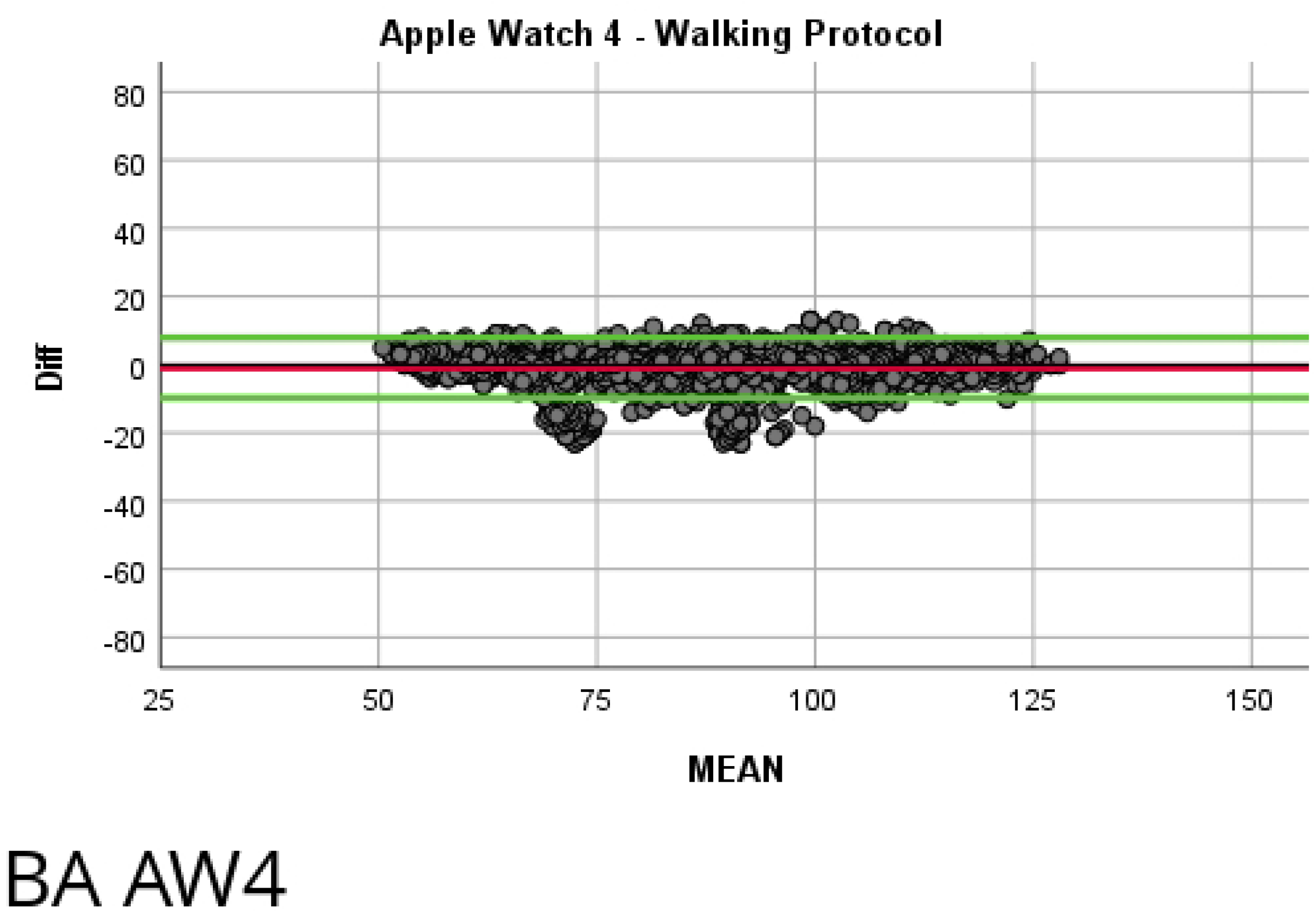
Bland-Altman Plot of all protocols for Garmin Forerunner 945. Mean bias of 1.60 with upper and lower limits of agreement of 20.469 and −17.269, respectively.

**Table 3.**
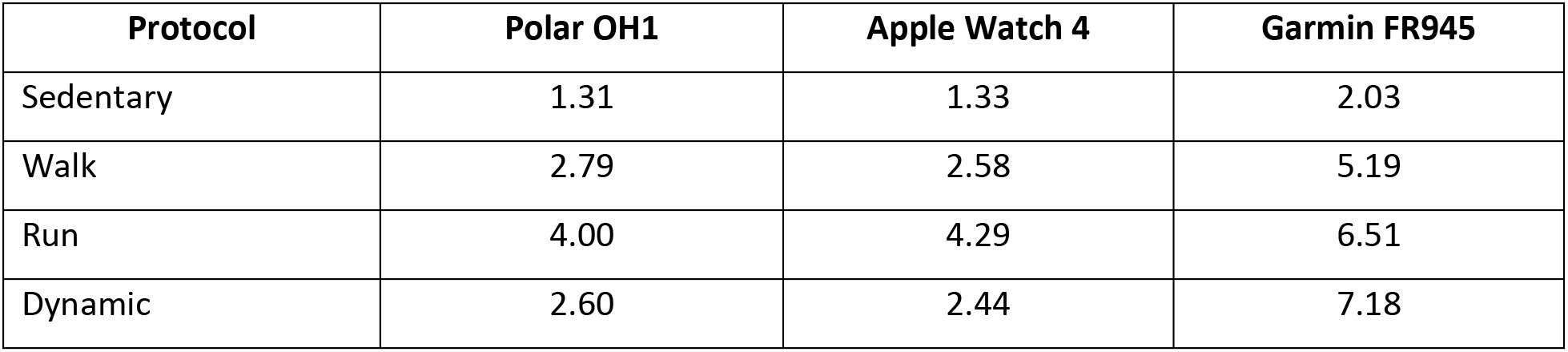
Mean Absolute Deviation (MAD)

| Protocol | Polar OH1 | Apple Watch 4 | Garmin FR945 |
| --- | --- | --- | --- |
| Sedentary | 1.31 | 1.33 | 2.03 |
| Walk | 2.79 | 2.58 | 5.19 |
| Run | 4.00 | 4.29 | 6.51 |
| Dynamic | 2.60 | 2.44 | 7.18 |

**Table 4.** Mean Absolute Percent Error (MAPE)

| Protocol | Polar OH1 | Apple Watch 4 | Garmin FR945 |
| --- | --- | --- | --- |
| Sedentary | 1.78% | 1.89% | 2.81% |
| Walk | 3.15% | 3.18% | 5.93% |
| Run | 3.43% | 3.51% | 5.25% |
| Dynamic | 2.05% | 1.95% | 5.47% |

#### Polar OH1

The OH1 resulted in a MAD between 1.31 (SED) and 4.00 (RUN) with a MAPE between 1.78% (SED) and 3.43% (RUN). The Bland Altman plot for the OH1 can be seen in Figure 1. The LoA for the OH1 ranged between −9.406 and 10.586 with a mean bias of .59. The ICC of the OH1 was .991 with 95% CI of .992 and .991.

#### Apple Watch 4

The AW4 produced a MAD between 1.33 (SED) and 4.29 (RUN). The MAPE for the AW4 ranged between 1.89% (SED) and 3.51% (RUN). The Bland Altman plot for the AW4 for all protocols (SED, WLK, RUN, DYN) can be seen in Figure 2. The overall Limits of Agreement (LoA) ranged from to −13.314 to 13.974 with a mean bias of .33. Intraclass Correlation (ICC) was high at .990 with a 95% Confidence Interval (CI) of .990 and .989, upper and lower, respectively. Pearson’s r was .990.

#### Garmin Forerunner 945

The FR945 results yielded a MAD between 2.03 (SED) and 7.18 (DYN) with a MAPE range between 2.81% (SED) and 5.93% (WLK). The Bland Altman plot for all protocols can be seen in Figure 3. The FR945 produced LoA between −17.269 and 20.469 with a mean bias of 1.6. The ICC was .967 with a 95% CI of .970 and .965. Pearson’s r was .969.

## Discussion

### ANSI/CTA Standards Validity

The principal findings of our study were that HR measurement via PPG technology in the Polar OH1, Apple Watch 4, and Garmin Forerunner 945 met the criteria to be considered valid by the ANSI/CTA standards. All three devices had a MAPE <10% while being evaluated across a broad subject group comprised of adequate representation across various skin tones, BMI levels, and sex.

Over the past two decades, wearable fitness devices have progressed in both use and functionality resulting in a broad range of options for consumers. A major advancement is the integration of PPG technology into the devices. By establishing the proper color of the light-emitting diode and refining proprietary algorithms, manufacturers can now provide end-users with myriad physiological information in a single device without the need for a chest strap. The devices used in this study all use similar PPG technology, primarily differing in only the number of diodes and the manufacturer’s unique algorithms.

The development of the ANSI/CTA standards for determining device validity defines a framework that generally allows for a more equitable and diverse application of the device characteristics to the total population. This study represents one of the first studies that has developed the study design in strict accordance with the ANSI/CTA standards. Subject selection was not random, but instead, individuals were specifically recruited to meet the minimum percent of subject group standards such that age, sex, skin tone and BMI were all adequately represented in the subject group. Additionally, exercise conditions were specifically designed to adhere to the standards, and subject input was utilized to appropriately set intensities across a very diverse group of subjects. Although specific analysis of appropriate intensity matching is beyond the scope of this research, visual analysis of the data suggests that all subjects performed each test condition in alignment with the information provided. Therefore, by implementing a strict study design and appropriately selecting subjects based on the prescribed framework, the results of this study can be broadly applied to the general population.

### Comparison with Previous Studies

Previous studies have attempted to determine validity for various PPG devices, although to our knowledge, this is the first study to strictly apply the ANSI/CTA standards to subject selection and study design. The devices in this study have been directly and indirectly studied in conjunction with other devices or using different methodologies. As the consumer electronics market is constantly progressing and new devices are introduced to consumers fairly frequently, direct device comparison is limited and requires inclusion of different versions or generations of the devices. Although device manufacturers have been researched extensively during the past 6 to 7 years, precise comparison between this study’s devices and previous research is very limited. Of the devices tested in this study, the OH1 has had been researched the most. This is most likely because the OH1 has stayed consistent during its lifetime whereas other products have had generational changes or complete updates to the product line. The original OH1 was released in 2017 with only one major upgrade to the OH1 Plus (allowed ANT+ communication). At the same time, Apple has released 4 different Apple Watches, and Garmin progressively released new watches in the Forerunner series with the FR945 being released in late 2019.

#### Polar OH1

Multiple studies have previously provided ample evidence of the validity of the OH1. Schubert et al. found the mean bias to be slightly higher than the current study (.59 versus .76) but a narrower LoA (−9.406 and 10.586 versus −3.83 to 5.35), but is limited in application as the study compared only a mean heart rate for a yoga session while also suffering from unbalanced subject sex selection (n=15, 3 males) with limited BMI and Fitzpatrick Scale variation (3) A more recent study found a lower mean bias (.27) and narrower LoA (−4.68 to 5.22) than the current study (4). Direct comparison is difficult as subjects the previous study noted all subjects held the handrail potentially decreasing any motion artifact, and the study also lacked any diversity with Fitzpatrick Scale and BMI. A 2019 study assessing different activities resulted in lower biases for walking and running (.18 versus .41 and .37 versus 1.28, respectively) but this study was biased towards males (n=70, males = 54), did not report BMI, and although it referenced skin tone, specific subject representation of skin tone levels was not reported (6). Additionally, A more recent study has further confirmed the validity of the OH1 in various activities, and across all activities found a higher mean bias (1 versus .59), a broader LoA (−20 to 19 versus −9.406 to 10.586) with a lower *r* (.957 versus .991) compared to the current study, but like other studies lacked subject information about skin tone and BMI (13).

#### Apple Watch 4

Apple regularly releases new products on an annual basis. As such, direct assessment of the AW4 is difficult, but evaluation of previous versions is available in the research. Dooley at al. evaluated the first-generation Apple Watch across a wide range of BMI and exercise intensities finding a higher MAPE for walking (5.60% versus 3.18%) and running (6.70% versus 3.51%) compared to the current study (7). Although the study utilized different treadmill walking and running intensities, the heart rate data was only recorded for a single time point and Fitzpatrick Scale was not recorded. In 2019 Hwang et al. researched the Apple Watch 2, revealing a much tighter LoA (−6.0 to 3.9 versus −13.314 to 13.974) but a slightly lower mean bias (−1.0 versus .330) than the current study, but Hwang used a model of tachycardia with electrical pacing so direct comparisons are difficult to determine (8). Nelson et al. released research concerning the Apple Watch 3 in 2019 (9), finding a higher mean bias (1.80 versus .330) and higher MAPE (5.86% versus 2.63%) than our study but Nelson’s study was a single-subject free-living design comparing different devices. Lastly, Duking et al. investigated the AW4 but the authors did not calculate key validity metrics (MAD, MAPE, ICC) with only a slightly lower *r* (.97 versus .984) available for comparison to the current study (10). Only one of these studies actively recruited subjects with skin tone variations, but the delineation was limited to white and non-white and ethnicity/race, not a skin tone scale (7).

#### Garmin Forerunner 945

FR945 validity data is lacking in the literature. The prior device-specific research that is available generally concerns the Forerunner 235 versus this study’s 945. In Dooley’s 2017 study, the Garmin Forerunner 235 had large deviations from the criterion HR with as high as MAPE of 24.38% (2.81% to 5.93% for the current study). In 2019, Stove et al. also completed research on Garmin Forerunner 235 validity revealing much lower ICC values (.480 to .905 versus .895 to .973) compared to the current study, but had a limited number of heart rate data points as data was only recorded once per minute (11).

### Device Differences

Although all devices tested were deemed valid, differences in MAD and MAPE for different devices did exist. Whether these differences are functionally important is determined by the consumer. In respect to the criterion, the OH1 and AW4 tended to have lower differences for MAD and MAPE values, as well as a higher ICC and narrower CI compared to the FR945. Similarly, the ICC and r were lower for the FR945 compared to the other two devices with the OH1 having a very slightly higher ICC and r than the AW4.

The reasons for these differences could be due to multiple factors. First, as previously mentioned, the devices all differ for functionality and intended use. Secondly, although each device was worn according to manufacturer’s specifications, devices differed in wristband/armband material and the size of the recording device. The FR945 and AW4 are both wrist-worn monitors but differing styles and materials of the wristbands resulted in slightly different fitment for the devices on individual subjects due to variation in wrist diameter. The OH1 had the smallest recording device and was secured to the lower arm via a fabric elastic band. Although the technology for the PPG light-emitting diode, appears to be similar between devices, individual devices variances between the number of diodes and spacing of diodes is visually apparent. The most likely reason for the differences, though, is the manufacturer-specific algorithm that converts the PPG raw data to heart rate information. Other differences in proprietary technology, such as the device specific hardware and software for recording and processing also presumably exist.

### Limitations

Although this study was conducted according to the current ANSI/CTA standards, certain limitations do exist. First, the subject group (n=20) is considered the minimum subject group size and minimum percentage for the specific parameters of BMI and Fitzpatrick Scale. Although adequate for ANSI/CTA standards, future studies should consider a larger subject group so that those two parameters can be more intricately analyzed within the total subject group. Additionally, the results of this study can only be applied to the specific devices and their corporation-specific algorithms to compute heart rate from PPG signals. As the technology continues to advance, it is plausible that the corporations will refine the algorithms in attempts to improve validity. Lastly, the ANSI/CTA standards place limitations on the subject group such that individuals with tattoos in the sensor location should not be included in the study due to presumed alterations in how the photosensor reads the reflection of the capillary beds. As it can be argued that tattoos on the arm and wrist have become popularized as of late, the validity of these devices cannot be confirmed in this subgroup.

## Conclusions

As consumers are consistently utilizing a variety of devices to track health metrics which rely on heart rate measurements, it is vital that the PPG recording technology and manufacturer proprietary algorithms properly represent the actual heart rate of the individual. As the end-consumer of these devices represents a wide range of subject characteristics, it is equally important that the devices correctly record heart rate across variations in age, skin tone, sex, and BMI. By utilizing the ANSI/CTA standards for heart rate recording devices, this and future studies can be more confident that the data recorded by the device can be utilized confidently by the majority of the population. In this study, one of the first to implement a study design in accordance with the ANSI/CTA standards, the Polar OH1, Apple Watch 4, and Garmin Forerunner 945 were all deemed valid in their measurement of heart rate. Consumers of various age, sex, body composition, and skin tone can be confident that the heart rate data presented to them is within a strict range for validity and represents their unique characteristics.

## Acknowledgements

The authors would like to acknowledge Phung Tran, Robert Tippett, Jr., Shreya Kulkarni for their assistance in data collection. Additionally, the authors would like to acknowledge Greg Dodd for his assistance with data alignment between the AW4 and H10. Lastly, the authors would also like to thank the Dr. David E. Martin Sport Science Research Fund and the Atlanta Track Club for their respective financial and logistical contributions to the study.

## Conflict of Interest

The authors state no conflict of interests.

## Abbreviations

ANSI/CTA: American National Standards Institute/Consumer Technology Association
AW4: Apple Watch 4
BPM: Beats per minute
GF945: Garmin Forerunner Multi-function watch
H10: Polar H10 Chest Strap Heart Rate Monitor (criterion)
ICC: Intraclass correlation
MAD: Mean Absolute Deviation
MAPE: Mean Absolute Percent Error
OH1: Polar OH1 Armband Heart Rate Monitor
PPG: Photoplethysmography

## Appendix 1.

### Polar OH1 Detailed Results

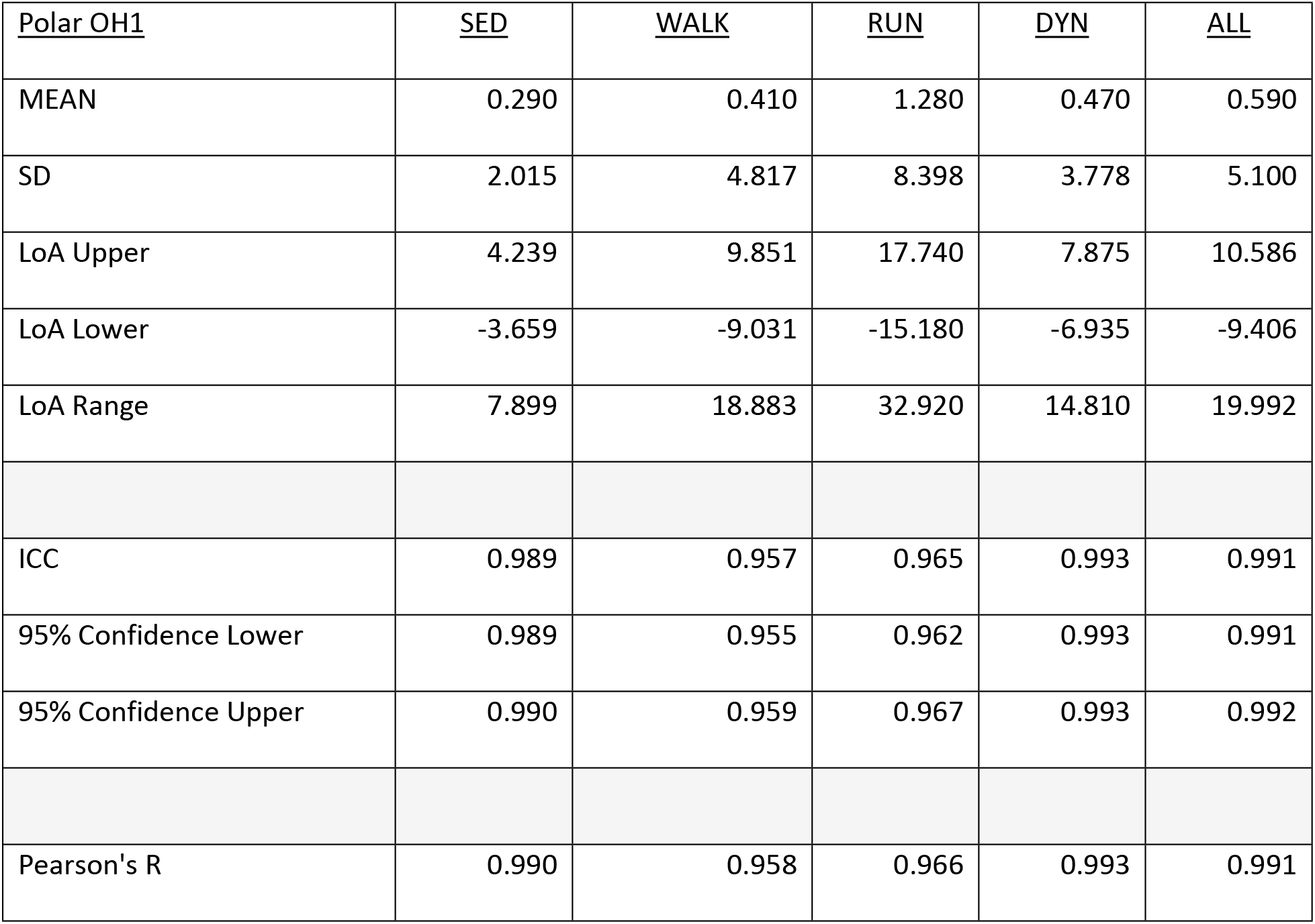

**Fig 4.**
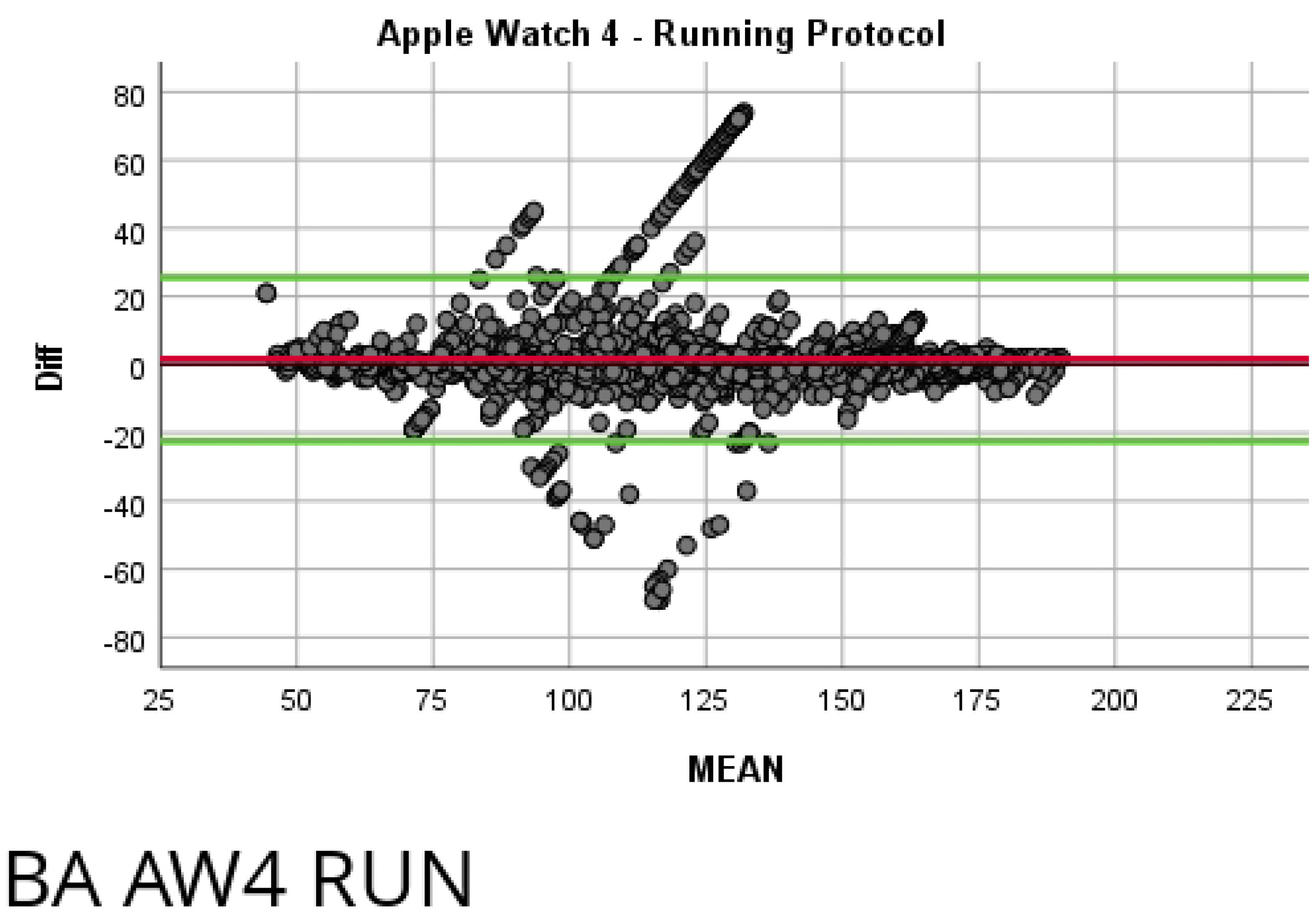
Bland-Altman Plot of Polar OH1 Sedentary Protocol. Mean bias of 2.015 with upper and lower limits of agreement of 4.239 and −3.659, respectively.

**Fig 5.**
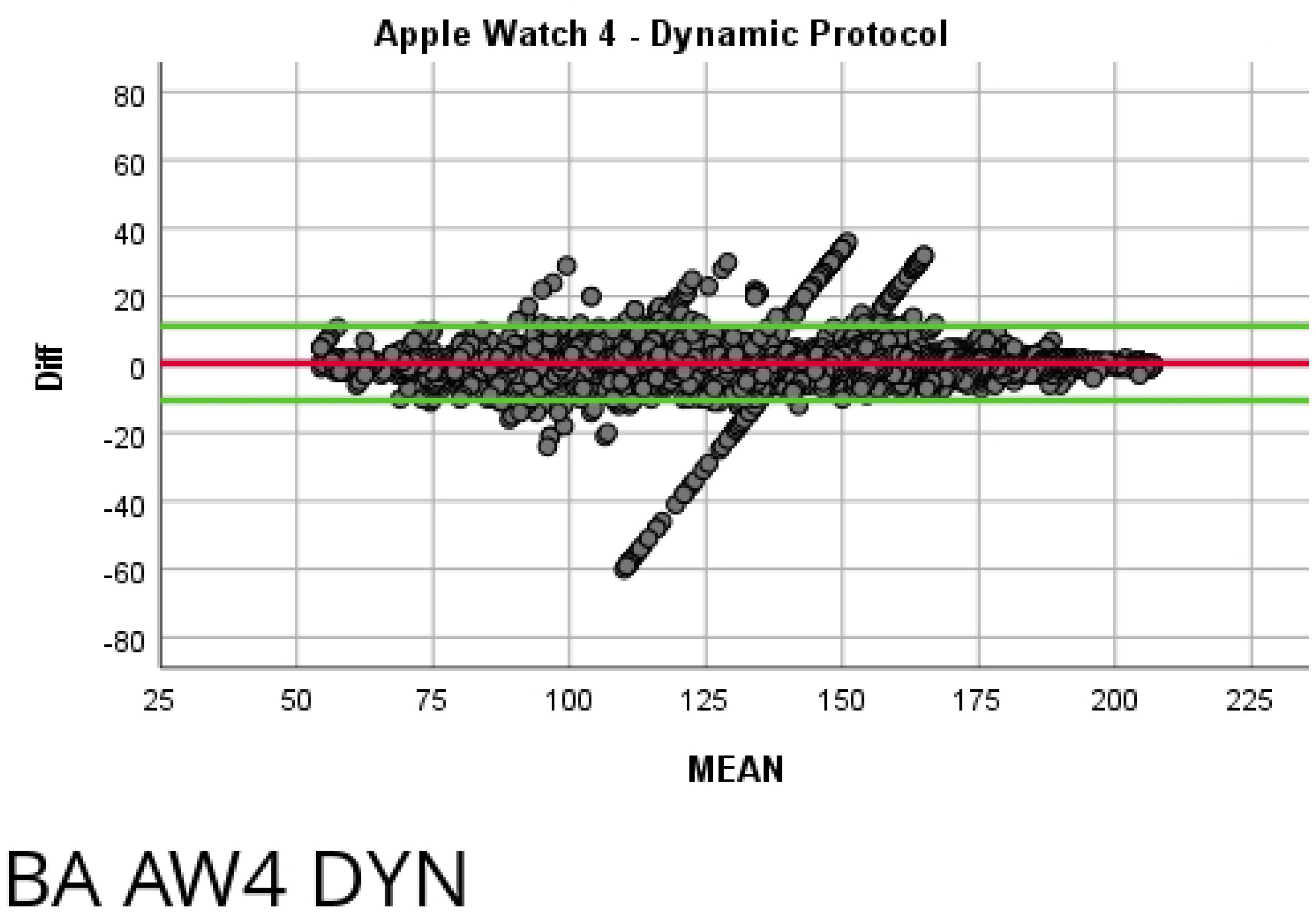
Bland-Altman Plot of Polar OH1 Walking Protocol. Mean bias of 4.817 with upper and lower limits of agreement of 9.851 and −9.031, respectively

**Fig 6.**
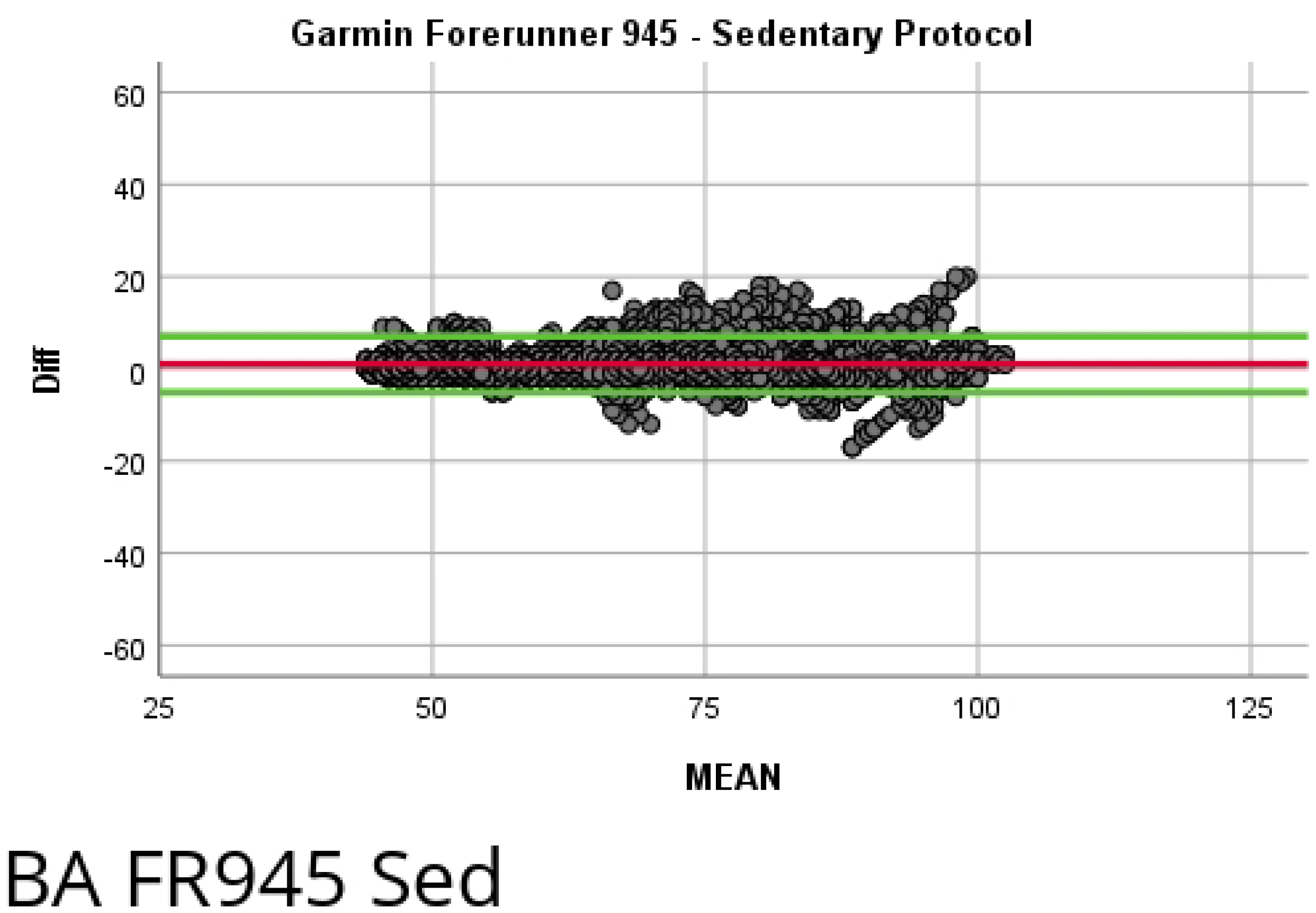
Bland-Altman Plot of Polar OH1 Running Protocol. Mean bias of 8.398 with upper and lower limits of agreement of 17.740 and −15.180 respectively.

**Fig 7.**
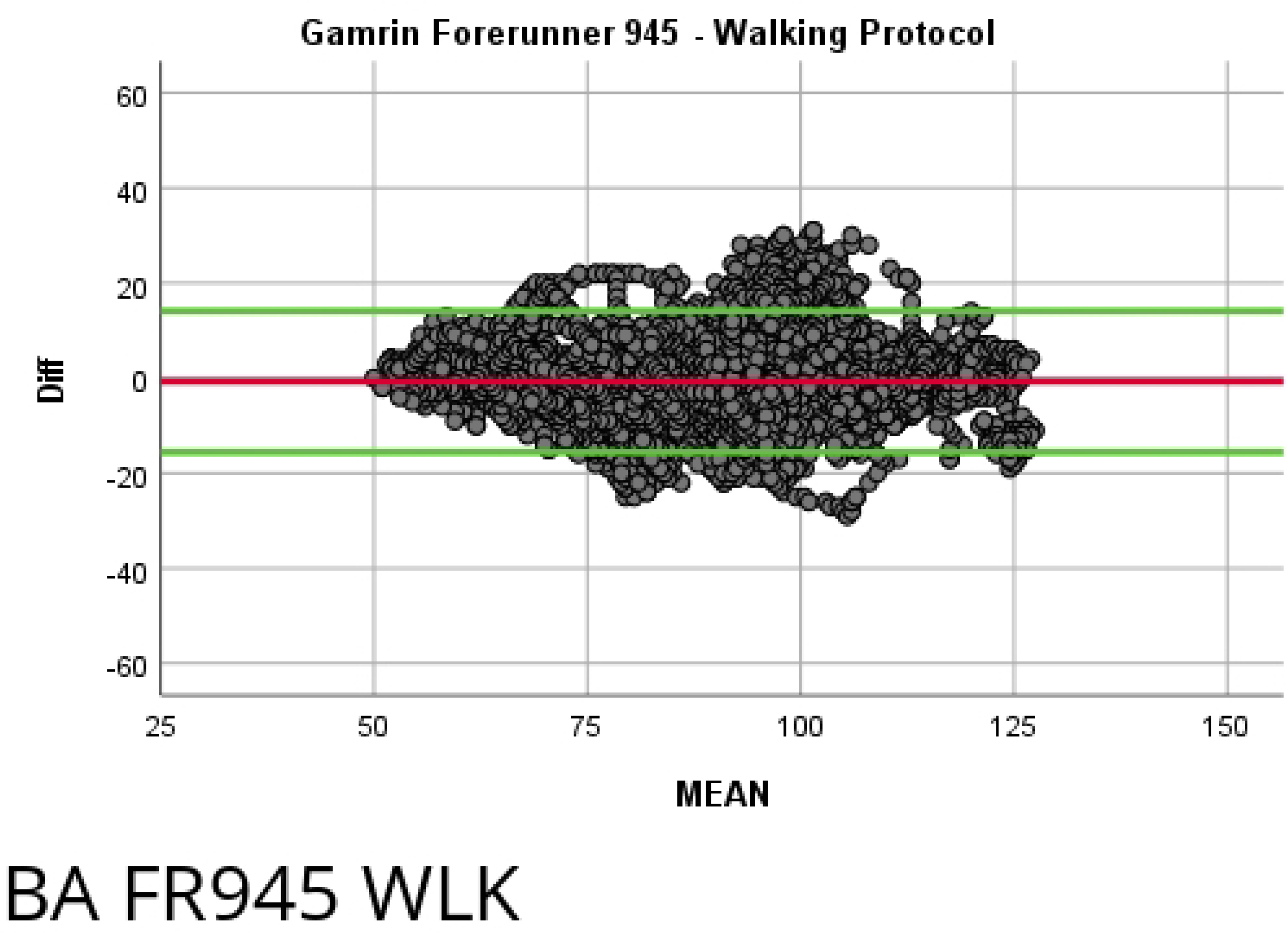
Bland-Altman Plot of Polar OH1 Dynamic Protocol. Mean bias of 3.778 with upper and lower limits of agreement of 7.875 and −6.935, respectively.

### Apple Watch 4 Detailed Results

| <u>Apple Watch 4</u> | <u>SED</u> | <u>WALK</u> | <u>RUN</u> | <u>DYN</u> | <u>ALL</u> |
| --- | --- | --- | --- | --- | --- |
| MEAN | 0.170 | -0.880 | 1.580 | 0.420 | 0.330 |
| SD | 2.020 | 4.500 | 12.257 | 5.589 | 6.961 |
| LoA Upper | 4.129 | 7.940 | 25.604 | 11.374 | 13.974 |
| LoA Lower | -3.789 | -9.700 | -22.444 | -10.534 | -13.314 |
| LoA Range | 7.918 | 17.640 | 48.047 | 21.909 | 27.287 |
| ICC | 0.990 | 0.960 | 0.926 | 0.985 | 0.984 |
| 95% Confidence Lower | 0.989 | 0.956 | 0.921 | 0.984 | 0.984 |
| 95% Confidence Upper | 0.990 | 0.965 | 0.931 | 0.985 | 0.984 |
| Pearson's R | 0.990 | 0.962 | 0.927 | 0.985 | 0.984 |

**Fig 8.**
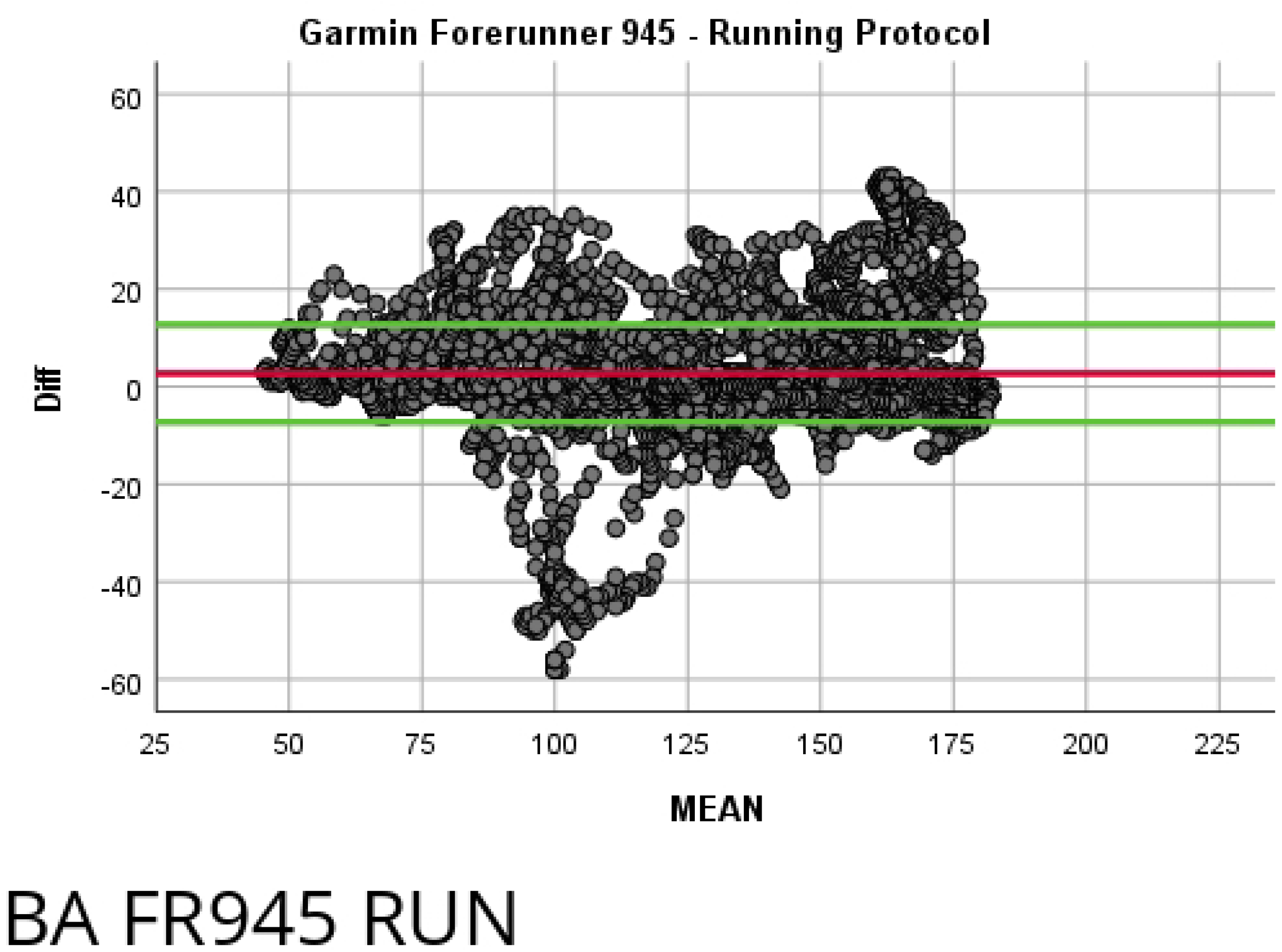
Bland-Altman Plot of Apple Watch 4 Sedentary Protocol. Mean bias of 2.020 with upper and lower limits of agreement of 4.129 and −3.789, respectively.

**Fig 9.**
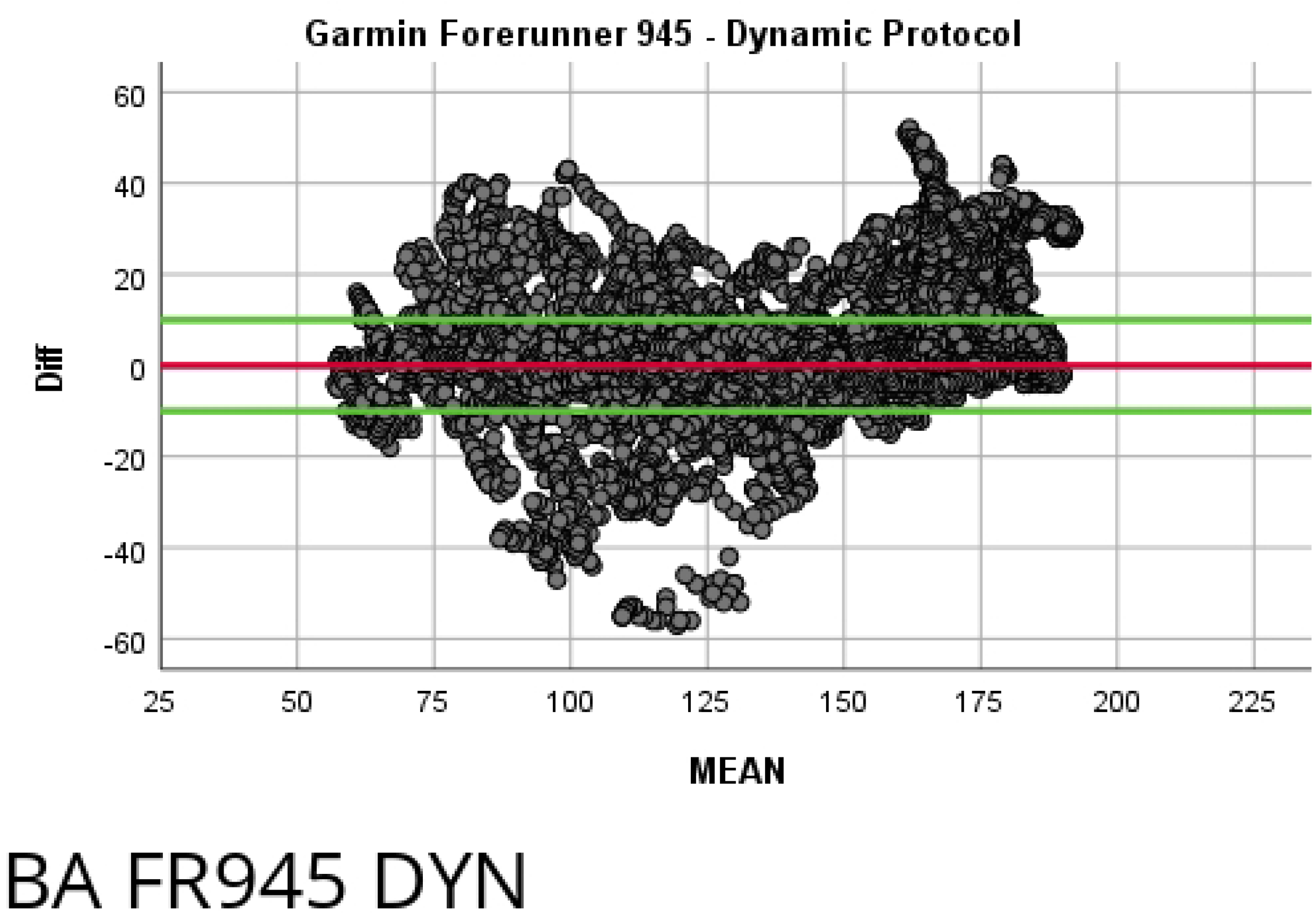
Bland-Altman Plot of Apple Watch 4 Walking Protocol. Mean bias of 4.500 with upper and lower limits of agreement of 7.940 and −9.700, respectively.

**Fig 10.**
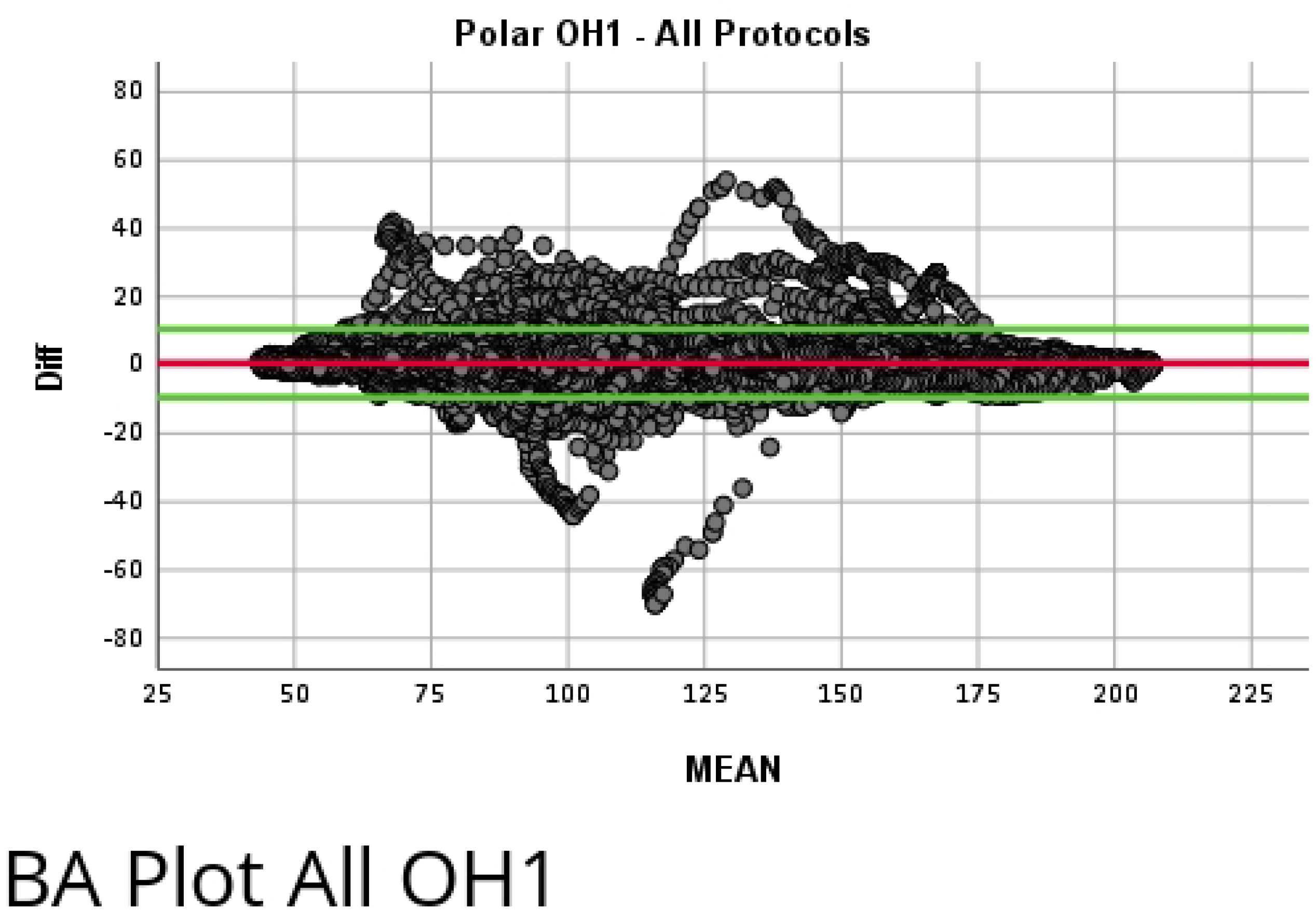
Bland-Altman Plot of Apple Watch 4 Running Protocol. Mean bias of 12.257 with upper and lower limits of agreement of 25.604 and −22.444, respectively.

**Fig 11.**
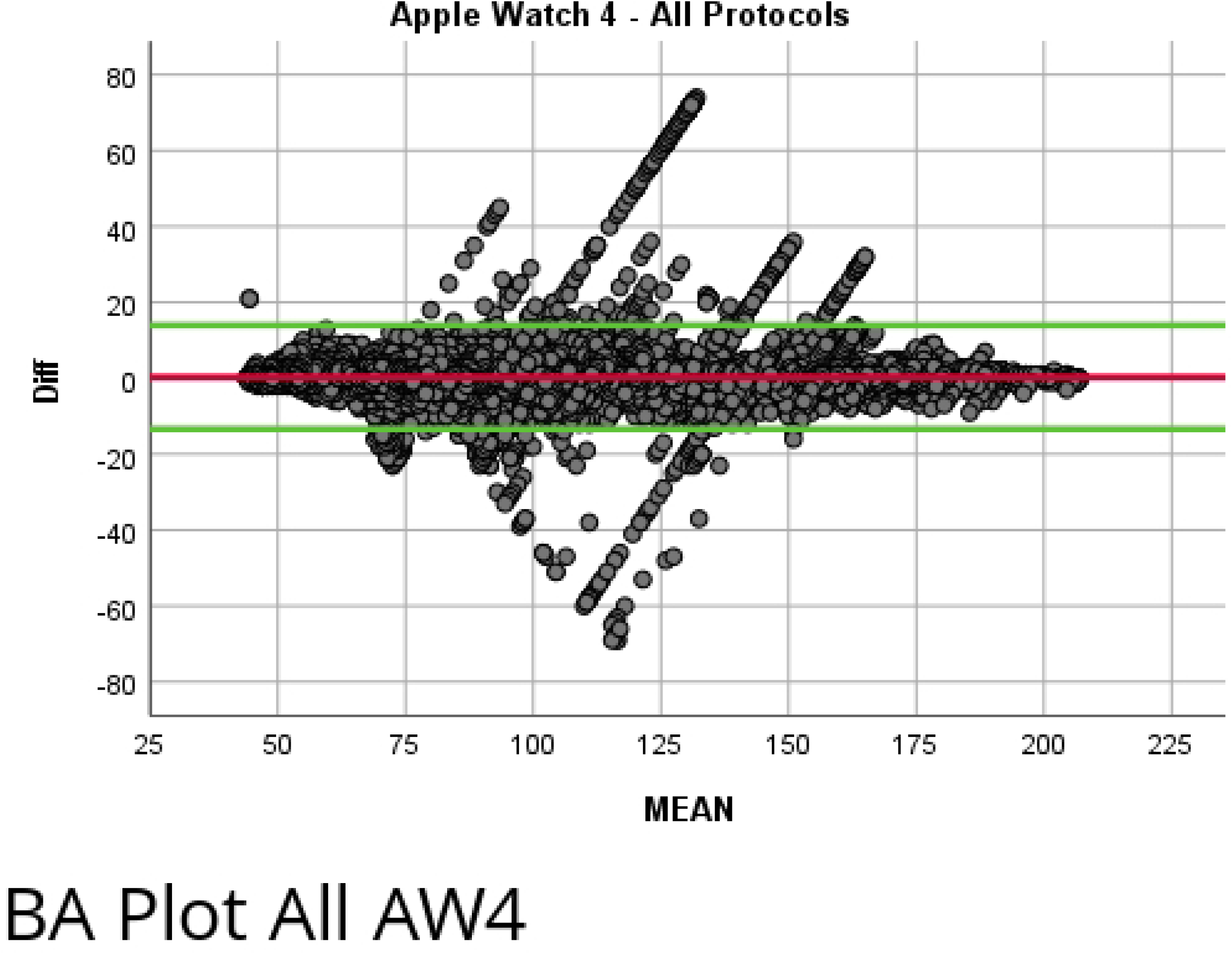
Bland-Altman Plot of Apple Watch 4 Dynamic Protocol. Mean bias of 5.589 with upper and lower limits of agreement of 11.374 and −10.534, respectively.

### Garmin Forerunner 945 Detailed Results

| <u>Garmin FR945</u> | <u>SED</u> | <u>WALK</u> | <u>RUN</u> | <u>DYN</u> | <u>ALL</u> |
| --- | --- | --- | --- | --- | --- |
| MEAN | 1.020 | -0.710 | 2.740 | 2.620 | 1.600 |
| SD | 3.106 | 7.590 | 11.158 | 11.760 | 9.627 |
| LoA Upper | 7.108 | 14.166 | 24.610 | 25.670 | 20.469 |
| LoA Lower | -5.068 | -15.586 | -19.130 | -20.430 | -17.269 |
| LoA Range | 12.176 | 29.753 | 43.739 | 46.099 | 37.738 |
| ICC | 0.973 | 0.895 | 0.934 | 0.922 | 0.967 |
| 95% Confidence Lower | 0.963 | 0.890 | 0.922 | 0.912 | 0.965 |
| 95% Confidence Upper | 0.979 | 0.900 | 0.943 | 0.931 | 0.970 |
| Pearson's R | 0.975 | 0.896 | 0.938 | 0.930 | 0.969 |

**Fig 12.**
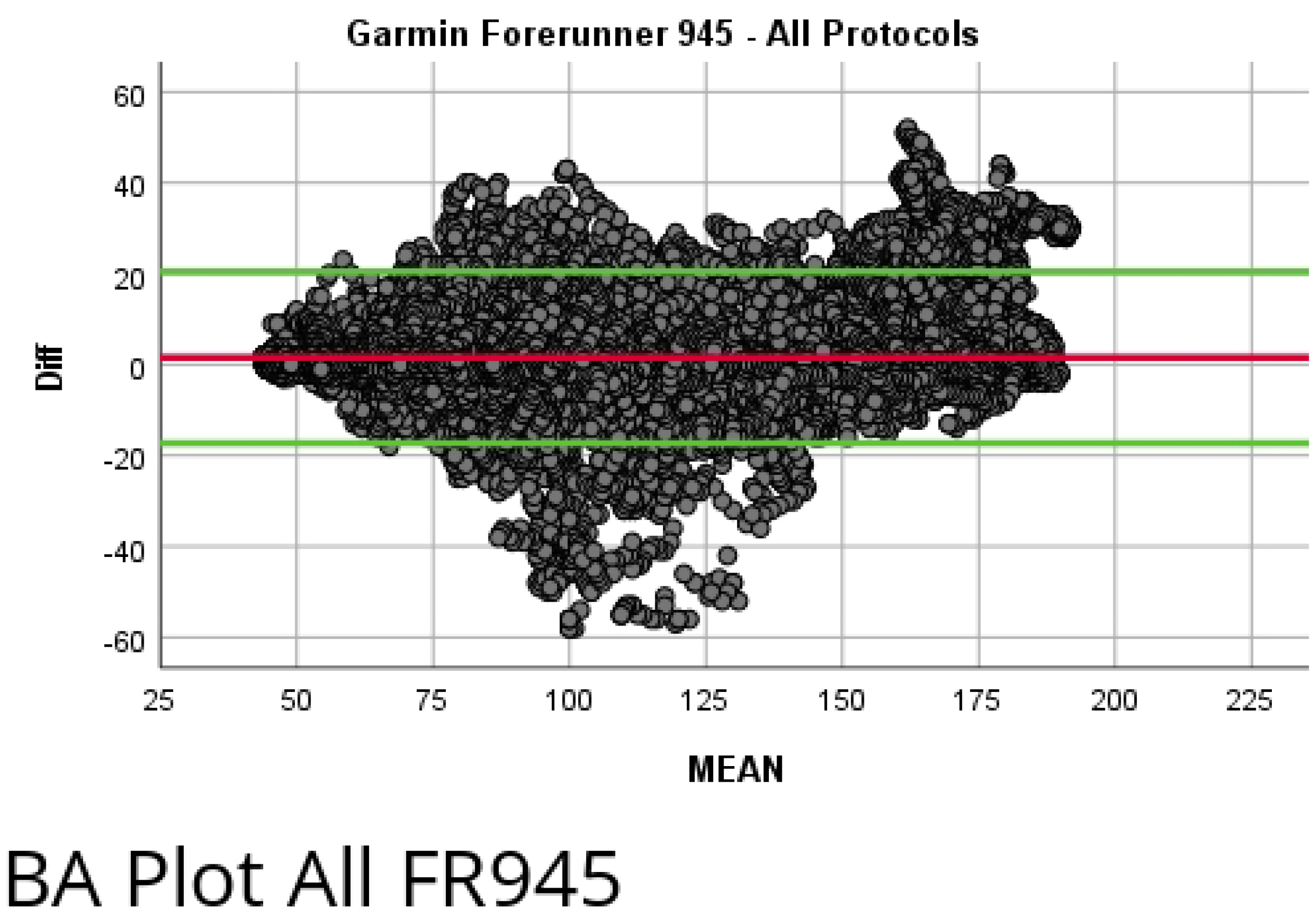
Bland-Altman Plot of Garmin Forerunner 945 Sedentary Protocol. Mean bias of 3.106 with upper and lower limits of agreement of 7.108 and −5.068, respectively.

**Fig 13.**
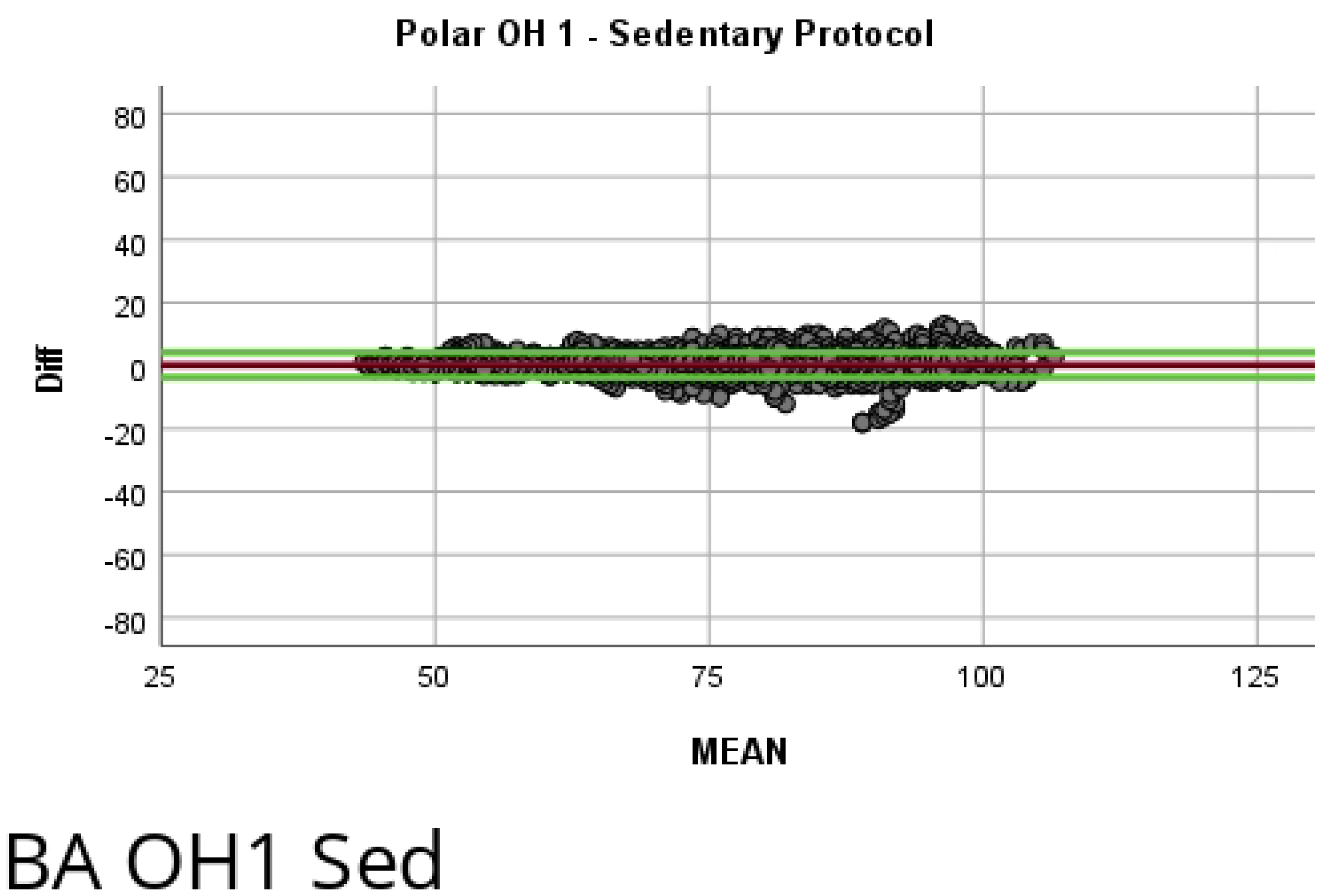
Bland-Altman Plot of Garmin Forerunner 945 Walking Protocol. Mean bias of 7.590 with upper and lower limits of agreement of 14.166 and −15.586, respectively.

**Fig 14.**
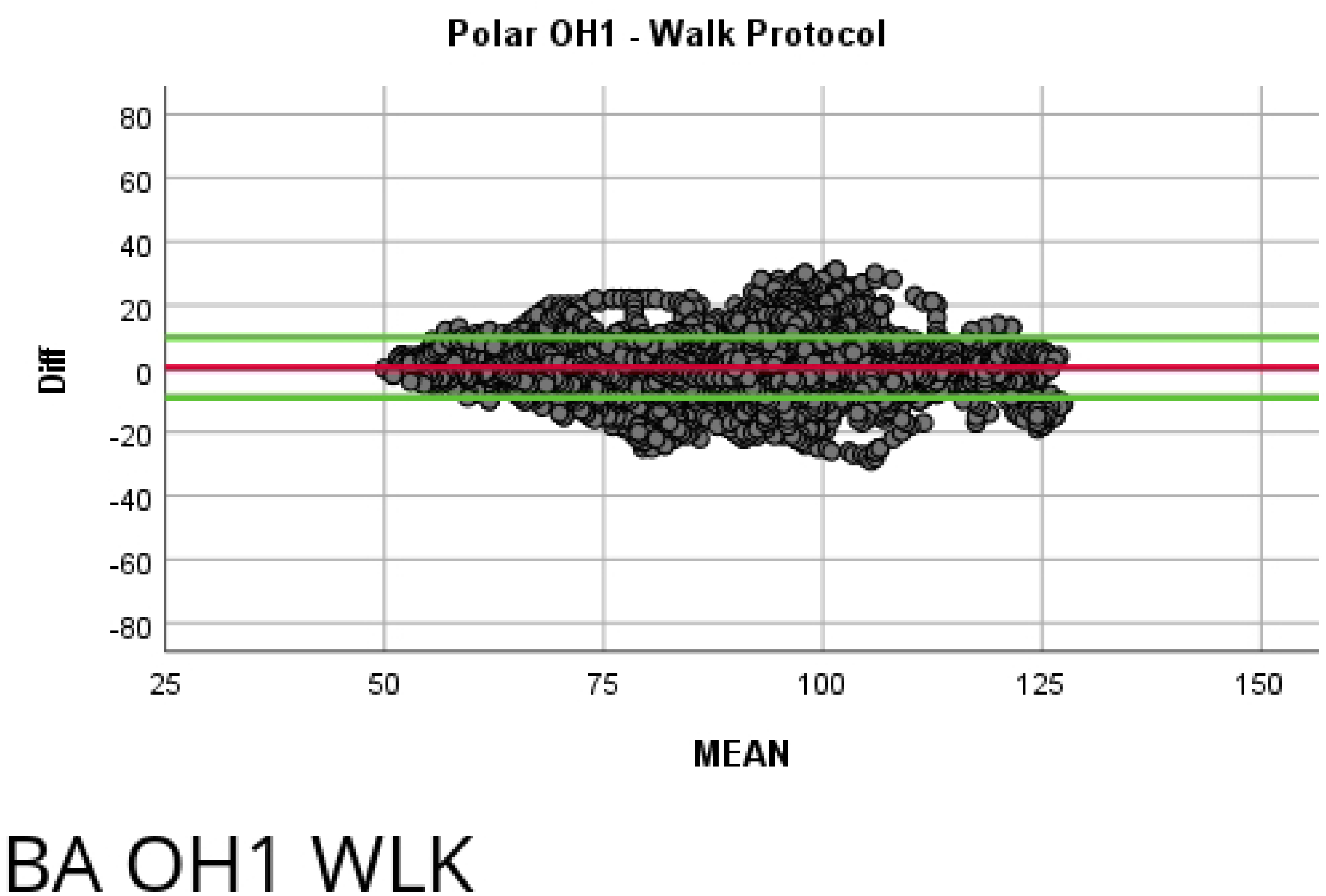
Bland-Altman Plot of Garmin Forerunner 945 Running Protocol. Mean bias of 11.158 with upper and lower limits of agreement of 24.610 and −19.130, respectively.

**Fig 15.**
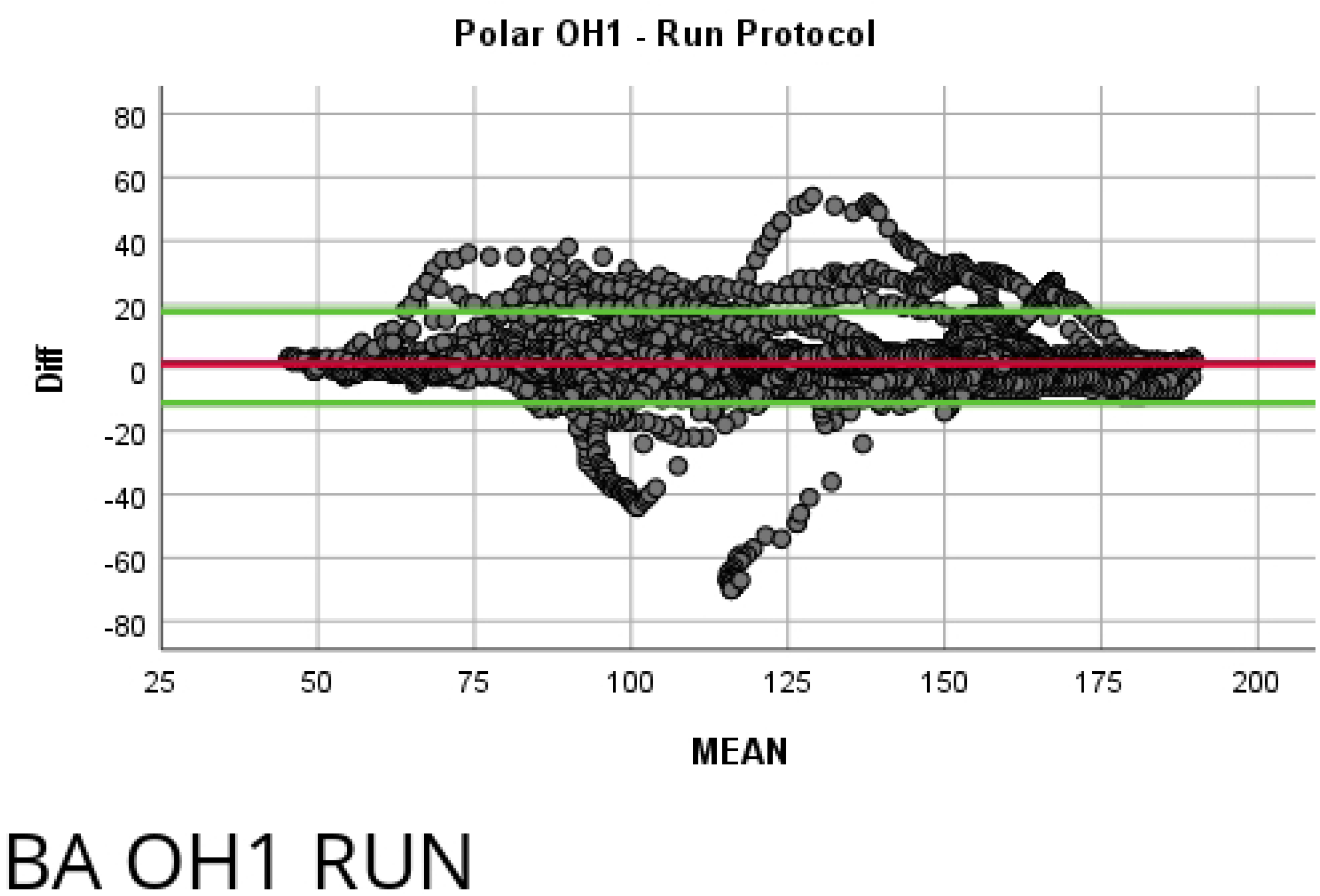
Bland-Altman Plot of Garmin Forerunner 945 Dynamic Protocol. Mean bias of 11.760 with upper and lower limits of agreement of 20.469 and −17.269, respectively.

## Notes

### Competing Interest Statement

The authors have declared no competing interest.

